# Comparative analysis of cattle breeds as satellite cell donors for cultured beef

**DOI:** 10.1101/2022.01.14.476358

**Authors:** Lea Melzener, Shijie Ding, Rui Hueber, Tobias Messmer, Guanghong Zhou, Mark J Post, Joshua E Flack

**Affiliations:** Mosa Meat B.V., Maastricht, The Netherlands; Department of Physiology, Maastricht University, Maastricht, The Netherlands; Lab of Meat Processing and Quality Control, College of Food Science and Technology, Nanjing Agricultural University, Nanjing, China

**Keywords:** Cultured meat, cattle breeds, satellite cells, proliferation, myogenic differentiation

## Abstract

**Background:** Cultured meat is a promising new field with the potential for considerable environmental and animal welfare benefits. One technological approach to cultured meat production utilises the proliferative and differentiative capacity of muscle-derived satellite cells (SCs) to produce large volumes of cultured muscle tissue from small biopsies of donor animals. Differing genotypes between cattle breeds lead to predictable phenotypic traits, resulting in breeds being favoured for their respective meat or milk production characteristics in the livestock industry. However, whilst these breeds show significant differences in muscle growth, it is unclear whether the physiological differences observed between them *in vivo* are reflected in differences in SC behaviour *in vitro*, particularly with respect to proliferation, differentiation and cellular longevity, and hence whether particular breeds might represent preferred SC donors for a cultured beef bioprocess.

**Results:** Comparing SCs isolated from five breeds (Belgian Blue, Holstein Friesian, Galloway, Limousin and Simmental), we found that the proliferation rates were largely unaffected by the donor breed. In contrast, potentially meaningful differences were observed in the kinetics and extent of myogenic differentiation. Furthermore, whilst differentiation dropped for all breeds with increasing population doublings (PDs), SCs from Belgian Blue and Limousin cattle showed significantly longer retention of differentiation capacity over long-term passaging.

**Conclusion:** SCs from all breeds were able to proliferate and differentiate, although Limousin and (particularly) Belgian Blue cattle, both breeds commonly used for traditional meat production, may represent preferred donors for cultured beef production.

## Introduction

Cultured meat (also referred to as ‘cell-based’ or ‘cultivated’ meat) is an emerging area of biotechnology that utilises the proliferation and differentiation of stem cells to produce mature, edible tissues for human consumption *in vitro* (1–3). These technologies are primarily motivated by sustainability issues associated with meat production, and have the potential to be vastly more resource efficient and animal-friendly (4).

Multiple approaches to cultured meat development are being explored, including the use of embryonic stem (ES) cells (1,3,5,6), induced pluripotent stem (iPS) cells (3), and immortalised cell lines. However, arguably the simplest approach utilises adult stem cells with the potential for myogenic differentiation, such as muscle satellite cells (SCs), the limited proliferative capacity of which necessitates repeated collection of tissue biopsies from donor animals (7). Bottlenecks in such a cultured meat bioprocess are thus the number of SCs isolated from a given mass of sample tissue, the rate and extent of myogenic differentiation, and, critically, the limits of proliferation that cells can undergo while maintaining the capacity for robust differentiation (8). Differences between donor animals with respect to these characteristics are therefore of crucial importance.

One particularly anticipated product is cultured beef, due to the extreme ecological impact of traditional cattle farming (9). Major genetic and phenotypic differences exist between cattle breeds, and particular traits (including maximum size and rate of weight gain) have been selectively bred for in order to optimise meat and/or milk production. For example, mutations in the *MSTN* (myostatin) locus are associated with a pronounced ‘double muscle’ phenotype in Belgian Blue and certain other breeds of cattle (10). Reduced or dysfunctional expression of myostatin leads to elevated expression of a myogenic transcriptional program during development (11,12). However, whilst some studies have hinted at differential trends in the behaviour of SCs isolated from different cattle breeds (13,14), it is still largely unclear to what extent the major physiological differences between breeds are reflected in the cell biology of their respective SCs *in vitro*.

In this study, we compared five cattle breeds traditionally used in different sectors of the livestock industry (outlined in Fig. 1a), with respect to the isolation, proliferation, differentiation and longevity of SCs. We use this knowledge to assess, for the first time, the suitability of these breeds as donor animals for cultured beef production.

**Figure 1:**
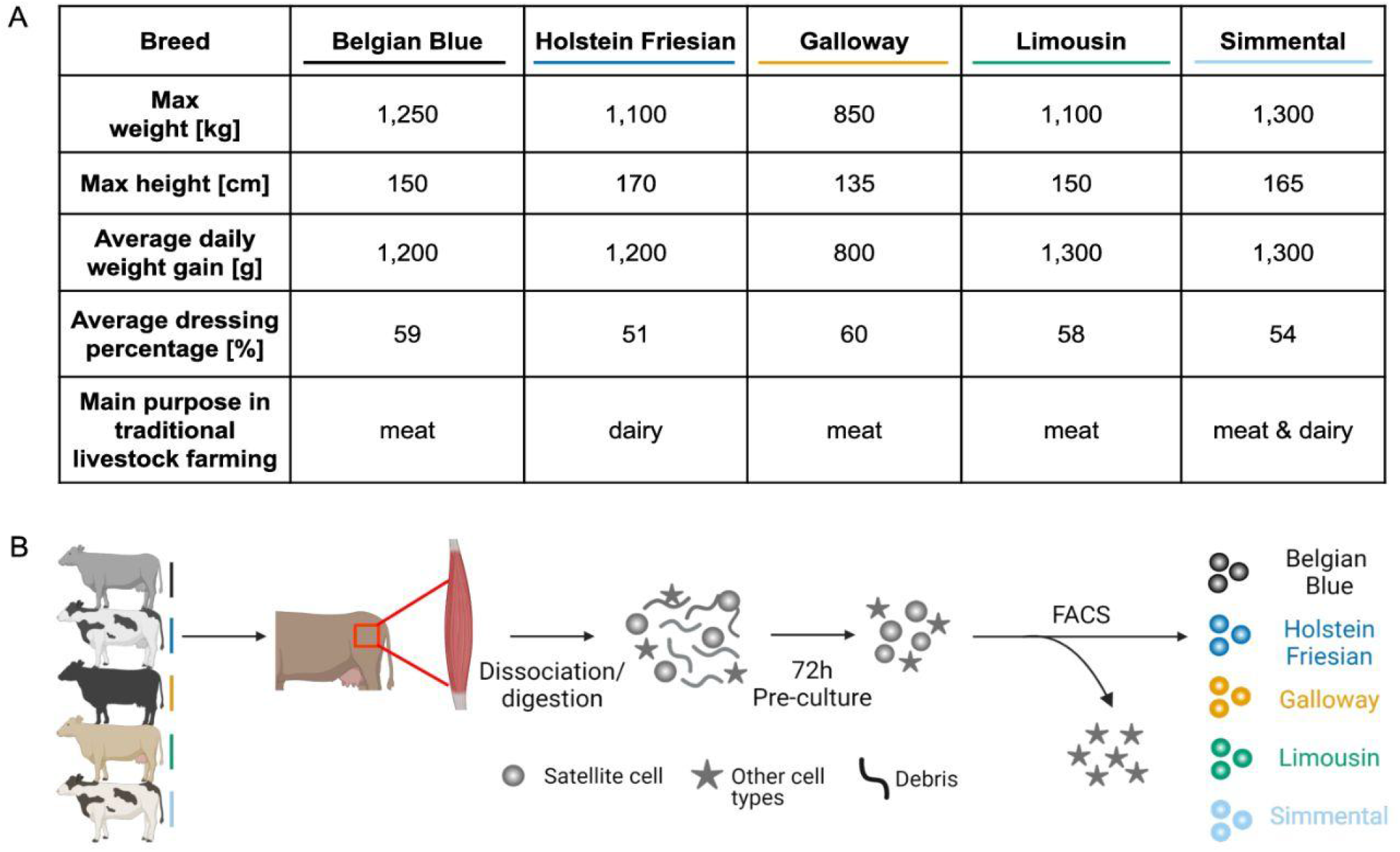
Comparative analysis of cattle breeds as satellite cells donors for cultured beef. **A**: Key phenotypic and agricultural characteristics of cattle breeds compared in this study (1–3). **B**: Overview of satellite cell harvesting and purification phase of the comparative study.

## Results

### Satellite cells can be purified from all cattle breed donors

In order to compare the suitability of cattle breeds as donors for cultured meat production, we first set out to isolate and purify SCs from semimembranosus muscle samples from heifers of each of these breeds (Fig. 1). We employed a fluorescence-activated cell sorting (FACS) strategy identical to that previously used for the isolation of SCs (Fig. 2a) (18). Whilst broadly similar, some differences were observable between breeds, including a reduction in the CD31/CD45+ population for the Belgian Blue sample. For all five breeds, a clear population of SCs (with a CD31-, CD45-, CD29+, CD56+ immunophenotype) was observable, although the size of this population varied slightly (from 6.9% in Limousin, to 16.5% in Holstein Friesian; Fig. 2a). Sorting this SC population yielded, for all breeds, a culture of mononuclear cells which adhered readily to collagen-coated tissue culture plastic (Fig. 2b). No obvious differences in cell morphology or clustering behaviour were observable by brightfield microscopy.

**Figure 2:**
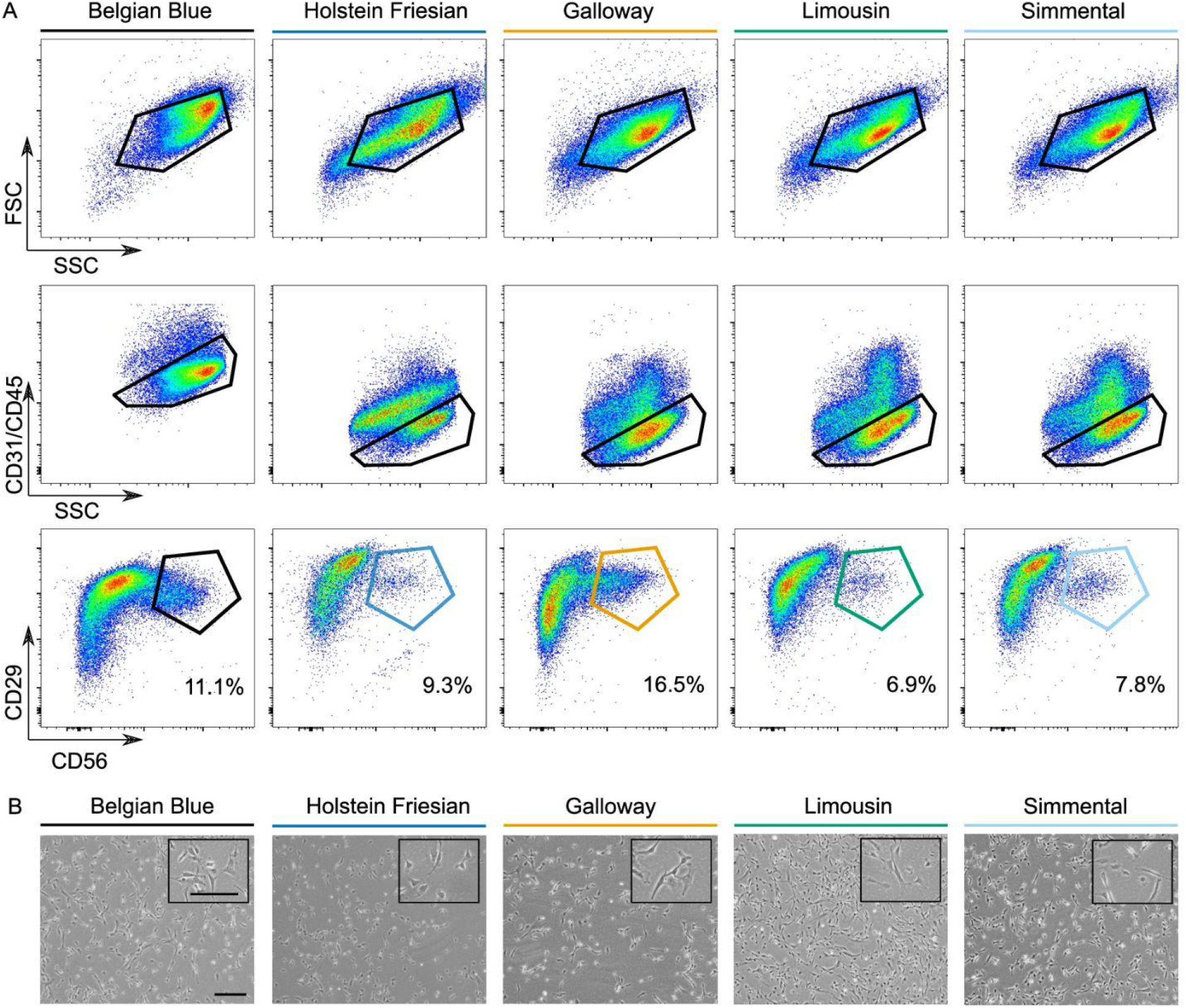
Identification and purification of satellite cells. **A:** Representative flow cytometry plots of unsorted bovine muscle cells from the five indicated cattle breeds, based on forward/side scatter (FSC/SSC respectively) and surface expression of CD31/CD45 (FITC), CD29 (APC) and CD56 (PE/Cy7). Coloured gates indicate fluorescence-activated cell sorting (FACS) strategy. Figures denote the percentage of satellite cells, as a proportion of the parent population. **B:** Brightfield microscopy images of sorted satellite cells from the indicated breeds. Scale bars = 20 μm.

### SCs vary in speed and extent of differentiation

We next wanted to compare the five cattle breeds in terms of the speed and extent of SC myogenic differentiation *in vitro*, both important parameters with respect to a cultured meat bioprocess. We induced differentiation of early passage SCs from each breed through serum-starvation (19), and followed the resultant changes at the microscopic, RNA and protein levels (Fig. 3) for 6 days. Differences in the kinetics of differentiation were more striking than in the maximum extent of differentiation reached. Whilst SCs from Belgian Blue and Limousin showed notably greater accumulation of myotubes after 3 days (Figs. 3a, b), this difference was reduced when comparing images of the SCs at their visual maximum extent of differentiation (Fig. 3c), suggesting that differentiation for the Holstein Friesian, Galloway and Simmental breeds may simply be slower. Indeed, whilst fusion index (the proportion of nuclei located inside desmin-positive myotubes (20)) was higher for Belgian Blue and Limousin at day 3 (Fig. 3d), morphological assessment of differentiation indicated that the difference between these and the other three breeds was significantly reduced between day 3 (Belgian Blue, p < 0.025; Limousin, p < 0.025) and day 6 (Belgian Blue and Limousin, not significant) of differentiation (Fig. 3e). We attempted to account for this variability in differentiation kinetics by comparing protein and RNA levels at the maximum extent of differentiation for each breed, and found that differences in expression of myogenic markers was nonetheless observable between the breeds. Although gene expression of key myogenic regulators and markers (21) was upregulated in all breeds, this was significantly higher for Belgian Blue SCs (Fig. 3f). The same trend was also observable in the respective upregulation of muscle-related protein levels (Fig. 3g).

**Figure 3:**
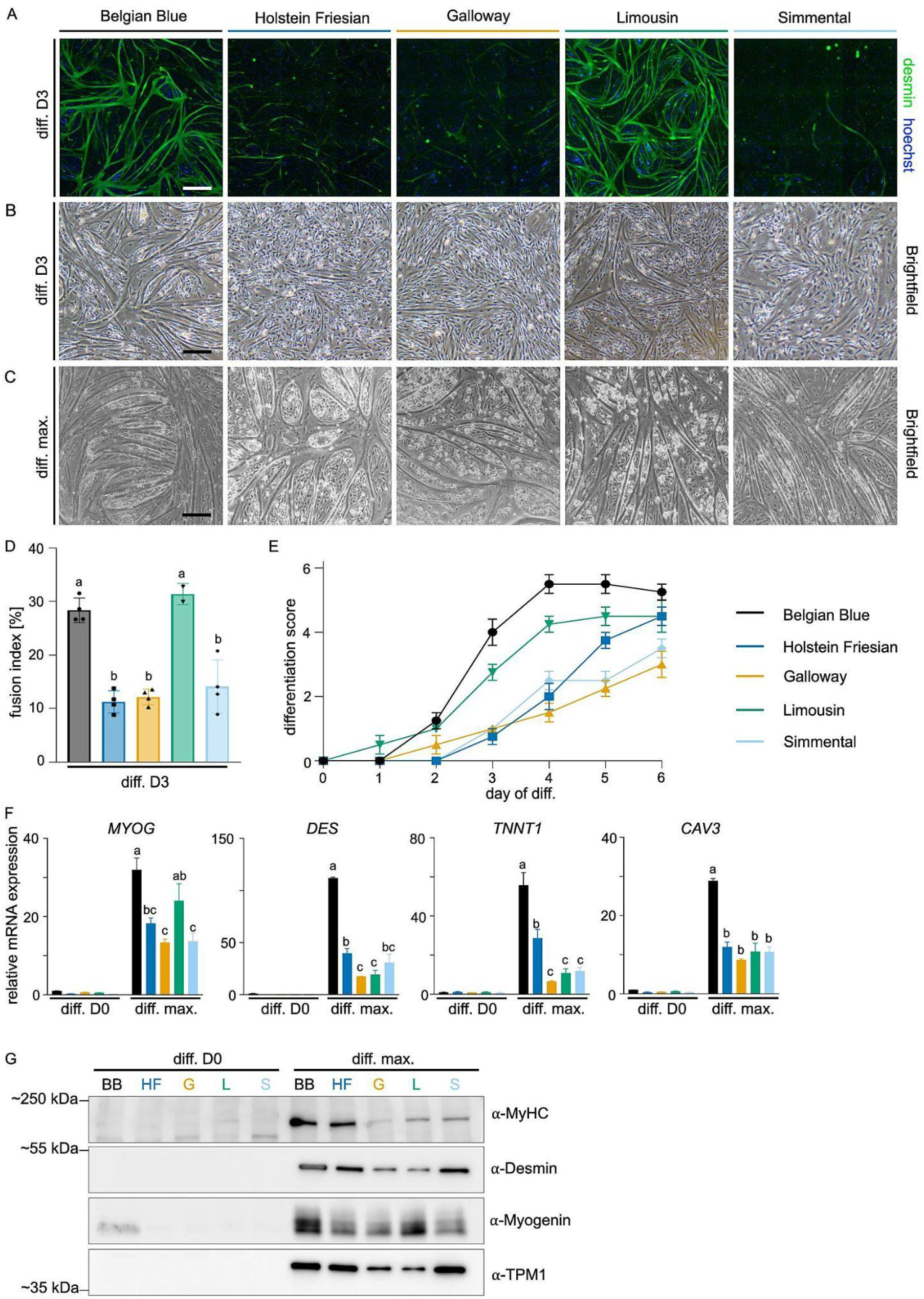
Comparative analysis of satellite cell differentiation. **A:** Representative immunofluorescence microscopy images of SCs from the five cattle breeds after 3 days of myogenic differentiation. Green = desmin, blue = Hoechst, scale bar = 20 μm. **B**: Brightfield microscopy images of Day 3 myogenic differentiation. Scale bar = 20 μm. **C:** Brightfield microscopy images on the day of maximum extent of myogenic differentiation. Scale bar = 20 μm. **D**: Quantified fusion indices of samples in A, presented as mean ± SEM (n = 4). Letters indicate clusters within which breeds do not differ significantly in a one-way ANOVA (P < 0.05). **E**: Differentiation scores based on visual assessment performed by 4 independent observers (κ = 0.49) of brightfield microscopy images on each day during differentiation. Data is presented in mean ± SEM (n = 4). Time (p < 0.0001), breed (p ≤ 0.0001) and the interaction of time and breed (p ≤ 0.0001) were all significant in a repeated measures test. **F**: Expression of genes specific for myogenic differentiation, as measured by RT-qPCR on Day 0 and on the day of maximum extent of differentiation for each breed. Gene expression was normalised against that in Belgian Blue, Day 0. Data is presented as mean ± SEM (n = 3). Time (p < 0.01), breed (p < 0.01) and the interaction of time and breed (p < 0.01) were all significant in two-way ANOVA. Post-hoc analysis: means not sharing a letter differ significantly in a Tukey’s multiple comparison test (P < 0.05). **G**: Western blot analysis of selected muscle-related proteins during myogenic differentiation measured on Day 0 and on the day of maximum differentiation for each breed.

### SCs differ between breeds with respect to longevity in culture

Although myogenic differentiation at early passage *in vitro* is a prerequisite for cultured meat production, maintenance of this differentiation capacity after an efficient, prolonged period of proliferation is also essential. Testing the long-term proliferative potential, we found that SCs from all five breeds were able to robustly proliferate for over 30 population doublings without a notable change in morphology (PDs; Figs. 4a, b). The observed proliferation rate declined slightly over the course of long-term passaging, although this did not differ significantly between breeds (Fig. 4b). Cycle analysis (as measured by the degree of EdU incorporation) showed a similar decreasing trend over passaging, although this was somewhat variable between breeds (Fig. 4c). In contrast, whilst the maximum extent of differentiation dropped significantly for all breeds over the course of long-term passaging, SCs from Belgian Blue (passage 9; p < 0.05) and Limousin (passage 9; p < 0.001) cattle demonstrated significantly better maintenance of differentiation potential from passages 6-9 (Fig. 4d), as evidenced by considerable myotube formation still occurring in these samples (Fig. 4e).

**Figure 4:**
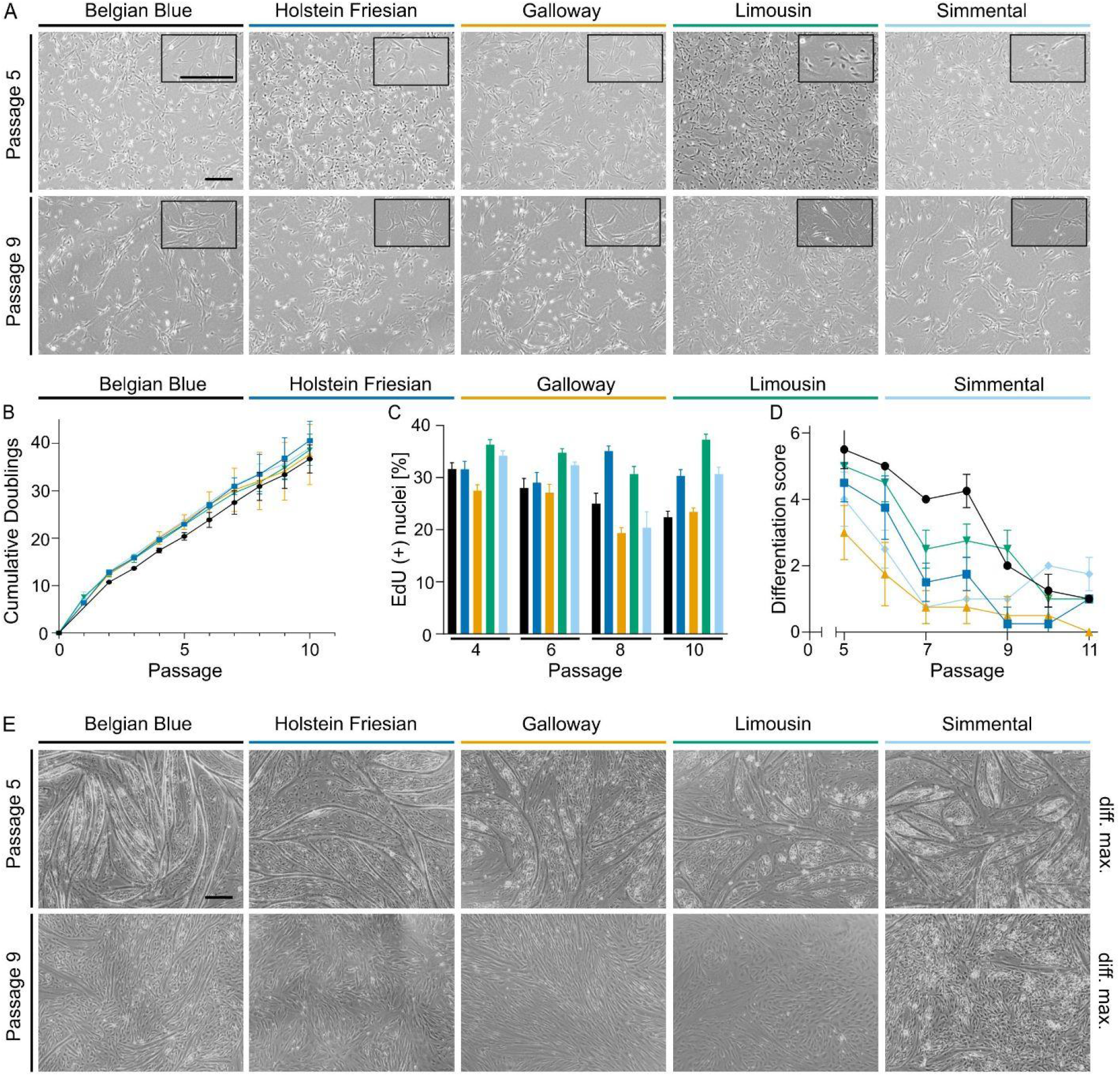
Long-term proliferation and differentiation. **A**: Representative brightfield microscopy images of satellite cell morphology with increasing passage number. Scale bar = 20 μm. **B**: Cumulative population doublings during long-term proliferation for the five indicated cattle breeds. Data is presented as mean ± SEM (n = 2). Time was significant (p < 0.0001), breed and and the interaction of time and breed were not significant in two-way ANOVA. **C**: Proportion of EdU positive cells during long-term proliferation in passage 4, 6, 8 & 10. Data is presented as mean ± SEM (n = 4). Time (p < 0.0001), breed (p < 0.0001), and the interaction of time and breed (p < 0.0001) were significant in two-way ANOVA. **D**: Maximum extent of differentiation over long-term proliferation, based on a visual differentiation scoring ranging from 0 to 6, performed by 4 independent observers (κ = 0.49). Data is presented as mean ± SEM (n = 4). Time (p < 0.0001), breed (p < 0.0001) and the interaction of time and breed (p < 0.0001) were all significant in two-way ANOVA. **E**: Representative brightfield microscopy images of myogenic differentiation with increasing passage number during long-term proliferation. Scale bar = 20 μm.

## Discussion

Design of an efficient cultured beef production process requires donor animals that yield SCs capable of robust proliferation and subsequent rapid differentiation into mature muscle fibers (1–3). In this study, we aimed to explore this issue by comparing the *in vitro* performance of SCs from matched donor cattle from five physiologically distinct breeds.

Using a previously identified FACS-based protocol (18), we were able to successfully purify SCs from all five cattle breeds (Fig. 2). The yield of SCs as a proportion of total cells was generally similar between all breeds, although the Holstein Friesian did display a moderately higher SC density. It has previously been observed that breeds with larger muscle fibres, such as Wagyu, contain relatively lower numbers of SCs per area of tissue than those with more modest muscle fibre diameters (22). It would be interesting to see whether such a mechanism also underlies the subtle differences observed in this study, although the relatively small differences between the sampled donors (combined with the exponential nature of the proliferative process, which places a greater importance on SC longevity) suggests that this is unlikely to be a major factor in determining optimal breeds for cultured beef production. Nevertheless, further optimisation of muscle sample harvesting will be required, particularly with respect to the switch to use of biopsied material (8). Differences in SC density between different muscles has previously been observed in pigs (23), and the proportion of fast- and slow-twitch fibres is also known to differ across these anatomical locations (24). Whilst intriguing, the potential relevance of these observations for cultured beef production has yet to be confirmed.

It is not fully clear how myogenic differentiation during the *in vitro* process of cultured meat production resembles early developmental myogenesis, as opposed to a wound healing process. Although differences in genotype between breeds did not manifest in obvious morphological differences between the isolated SCs, more notable differences in terms of myogenic differentiation were observed (Fig. 3). While all five breeds underwent differentiation, as measured at the RNA, protein and microscopic levels, the rate of this differentiation was markedly higher for Belgian Blue SCs. These cattle carry a well-described deletion in the *MSTN* (myostatin) gene, encoding a secreted TGFβ-family inhibitory myokine (10,25). The resultant loss of myogenic suppression leads to increased expression of transcription factors, including *MYOD1*, relative to other breeds (26), corroborating our observations. Interestingly, SCs derived from Limousin also showed rapid fusion, concomitant with elevated expression of *MYOG*, but not other muscle-related genes. Whilst Limousin and other beef breeds lack an obvious genotypic variant (in contrast to Belgian Blue), strong selection for meat specific traits has nonetheless resulted in significant polymorphism in the *MSTN* gene (27,28). Furthermore, it is likely that subtle changes in the regulation of underlying signalling pathways, such as IGF, are also involved (29). Recent work has additionally identified differential microRNA expression between SCs of different cattle breeds (14).

Furthermore, similar differences were also observed with respect to the longevity of the SCs in culture (in other words, the number of population doublings cells could undergo while maintaining robust differentiation capacity; Fig. 4). Whereas proliferation rates were largely similar between breeds, Belgian Blue- and Limousin-derived SCs showed notably improved differentiation after 8-9 passages. These differences in cellular longevity likely relate to alterations in cellular signalling (perhaps of the p38/MAPK pathway) between breeds, and hence differential responses to previously described chemical modulators of cell ageing might be expected (15).

The correlation between the early-passage differentiation and longevity of Belgian Blue (and to a lesser extent, Limousin) SCs suggests that increased sensitivity to activation of the myogenic transcriptional program might be a general feature of optimal donor breeds for cultured beef production. This will require validation in upscaled proliferation and differentiation systems, as opposed to the 2D model systems employed here (30). It also remains to be determined if different breeds show differential sensitivity to myogenic inducers in a serum-free setting, rather than the serum-starvation method used here. Furthermore, accurate mimicry of traditional meat requires the inclusion of fat tissue, which contributes significantly to taste and texture (31). Myostatin dysregulation is known to also impact adipogenesis (32), and it is not clear whether optimal SC donors would also represent good donors of adipogenic stem cells, such as fibro-adipogenic progenitors (FAPs) (8).

Whilst we have focussed on cattle breed in this study, age (33), sex and husbandry conditions (34,35) are also highly relevant factors that must be investigated for their impact on donor selection for cultured beef production. Although further work remains, this study nevertheless highlights the importance of careful donor animal selection and indicates that, to at least some extent, physiological differences between animals can be reflected in cell biological variation. In this instance, Limousin and (particularly) Belgian Blue cattle, breeds traditionally used for meat production, may represent optimal SC donors for cultured beef production due to their rapid and robust myogenic differentiation. Further characterisation of donor genomics and its correlation with *in vitro* traits will help to select for optimal donors in future, inform the selective breeding of desirable cultured beef related traits, and provide insights into donor selection for other types of cultured meat.

## Methods

### Animal care and welfare

Satellite cells were isolated from a heifer of each of five different cattle breeds. Housing conditions of the donor animals were similar for the five breeds. All were kept in a running pen with *ad libitum* access to a daily ration of a standard fattening feed. Heifers were slaughtered at an age of 30 ± 12.6 months. Holstein Friesian, Galloway, Limousin and Simmental cattle were kept and slaughtered in Germany, while Belgian Blue cattle were kept and slaughtered in the Netherlands, in accordance with national guidelines on animal tissue handling.

### Cell isolation

Immediately post-slaughter, samples were taken from the semimembranosus muscle and transported on ice. Muscle-derived cells were isolated using the method previously described (18). Briefly, small bundles of muscle fibres were collected and digested (1 h, 37°C) in the digestion buffer (Table S1). Repeated centrifugation was followed by filtering steps with 100 μm cell strainers. Cell pellets were incubated with erythrocyte lysis buffer (ACK; 1 min, room temperature), resuspended in growth medium (GM; Table S1), and filtered through 40 μm cell strainers. Cells were cryopreserved in freezing medium (Table S1), prior to long-term storage in liquid nitrogen.

### Fluorescence activated cell sorting (FACS)

Prior to FACS, unsorted cells were cultured for 60 hours on bovine collagen type I (Sigma, Cat# C2124; 2.5 μg/cm^2^) coated flasks in a humidified incubator (5% CO_2_, 37°C). Cells were sorted using antibodies as previously described (18) on a FACSAria cell sorter (BD). SCs were sorted by gating for the CD31/CD45-, CD29+/CD56+ population.

### Cell culture

Sorted SCs were cultured on collagen coated flasks in GM. Cells were seeded at 2,000 cells/cm^2^ and passaged every 3 to 4 days. For myogenic differentiation, SCs were seeded at 30,000 cells/cm^2^ on tissue culture plates coated with 0.5% Matrigel (BD Biosciences). Differentiation was induced at a confluency of approximately 90% by treating the cells with differentiation medium (DM; Table S1). Brightfield images were captured using a Nikon benchtop microscope (Eclipse TS100-F) and camera (DS-L3) system.

### Immunofluorescence microscopy

SCs were fixed with 4% paraformaldehyde (PFA), permeabilised with 0.5% Triton X-100, and blocked in 5% bovine serum albumin (BSA). Cells were stained with α-desmin antibody (D1033, Sigma Aldrich), secondary antibody (AF488 α-mouse; A11001, Invitrogen) and Hoechst 33342 (Thermo Scientific), and imaged using an ImageXpress Pico Automated Cell Imaging System (Molecular Devices).

### Brightfield microscopy differentiation image analysis

For each breed and passage, brightfield microscopy images were captured daily to create a 6 day differentiation time course. Images were scored (after removal of metadata and randomisation of ordering) by four independent observers based on a scoring system ranging from 0 (no differentiation) to 6 (high proportion of myotubes), as presented in Fig. S3.

### RT-qPCR

RNA was harvested from tissue culture samples by addition of TRK lysis buffer, and purified using the Omega MicroElute Total RNA Kit (Omega Bio-tek). RNA concentrations were determined by nanospectrometry, and reverse-transcribed using the iScript cDNA synthesis kit (Bio-Rad). Quantitative real-time PCR (RT-qPCR) was performed using iQ SYBR Green Supermix (Bio-Rad), with primer pairs shown in Table S2.

### Western blotting

Cells were lysed on ice using RIPA Lysis Buffer (sc-24948, Santa Cruz Biotechnology). Protein samples were boiled in Laemmli buffer (5 min), separated by SDS-Page and transferred to PVDF membranes by electrophoresis. Equal protein loading was confirmed by full-protein staining using Revert 700 Total protein stain (LI-COR; 926-11015). Blots were stained with primary antibodies against desmin (D1033, Sigma Aldrich), myogenin (sc-52903, Santa Cruz Biotechnology), α-tropomyosin (ab133292, Abcam), myosin IIα (M8421, Sigma Aldrich), and respective secondary antibodies α-mouse-HRP (ab6721, Abcam) or α-rabbit-HRP (P0447, Dako). Blots were developed using SuperSignal West Femto Maximum Sensitivity Substrate (Thermo Scientific) and visualised on an Azure 600 chemiluminescence imager (Azure biosystems).

### EdU-labeling

The Click-iT EdU Cell Proliferation Kit (C10337, Thermo Fisher) was used to assess cell cycle state. SCs were seeded at 2,000 cells/cm^2^ on collagen coated 96-well plates and cultured for 24 h, followed by incubation with 5-ethynyl-2’-deoxyuridine (EdU; 1 h). Cells were fixed with 4% PFA, permeabilised and stained according to the manufacturer’s instructions. Images were captured using the ImageXpress Pico Automated Cell Imaging System (Molecular Devices).

### Statistics

Statistical analyses were performed using Prism 9.1.0 (GraphPad). To compare the differences in differentiation over time of the cells isolated from the five different breeds, one-way ANOVA was performed (Fig. 3d). Repeated Measures was performed to compare the effect of beed, time and the interaction between these two parameters, when the samples were paired (Fig. 3e). Two-way ANOVA was used to distinguish the effects of the breeds and time respectively as well as the interaction of the two parameters when the samples were unpaired (Figs. 4a-c). Tukey’s multiple comparisons test was used as a post hoc analysis to demonstrate significance of differences between all groups (Figs. 3d-f; 4d).

## Author contributions

LM, SD, RH, TM and JEF performed experiments and analysis. GZ, MJP and JEF supervised the study. LM, MJP and JEF wrote the manuscript with input from all authors.

## Acknowledgements

We would like to thank Will Hart (Opvia Ltd) for software facilitating the scoring of myogenic differentiation images. We would also like to thank Karl Schellander, Christiane Neuhoff, Michael Hölker, and the staff at the Department of Animal Breeding and Husbandry, Institute of Animal Science, University of Bonn for facilitating sampling of the cattle breeds.

## Declared conflicts of interest

LM, RH, TM and JEF are employees of Mosa Meat B.V. SD is co-founder of Joes Future Food. MJP is co-founder and stakeholder of Mosa Meat B.V. Study was funded by Mosa Meat B.V.

