## Supplementary Material for "Comparative analysis of cattle breeds as satellite cell donors for cultured beef"

**Table S1: Media and buffer compositions.**

| Component | Concentration |
|---|---|
| **Digestion Buffer** | |
| DMEM (Invitrogen, Cat# 41966-29) | |
| Collagenase 1 (Worthington, Cat# CLS-1) | 0.2% |
| Penicillin/Streptomycin/Amphotericin (PSA; Lonza, Cat# 17-745E) | 1% |
| **Growth Medium (GM)** | |
| Ham's F-10 (Gibco, Cat# 31550-023) | |
| Heat-inactivated fetal bovine serum (FBS; Gibco, Cat# 10500064) | 20% |
| PSA (Lonza, Cat# 17-745E) | 1% |
| Recombinant human bFGF (R&D systems, Cat# 233-FB) | 5 ng/ml |
| **Freezing medium** | |
| Heat-inactivated FBS (Gibco, Cat# 10500064) | |
| DMSO | 10% |
| **Differentiation Medium (DM)** | |
| DMEM (Invitrogen, Cat# 41966-29) | |
| Heat-inactivated FBS (Gibco, Cat# 10500064) | 2% |
| PSA (Lonza, Cat# 17-745E) | 1% |

**Table S2: RT-qPCR primers.**

| Gene | Primer sequence | |
|---|---|---|
| *UXT* | Fwd: | 5'-GAGCAGTCTCCTCACAGAGCTC |
| | Rev: | 5'-AGCAACATGTGGATATGGGCCT |
| *L19* | Fwd: | 5'-TCGAATGCCCGAGAAGGTAAC |
| | Rev: | 5'-CTGTGATACATGTGGCGGTC |
| *CAV3* | Fwd: | 5'-GATCGATCTGGTGAACCGGG |
| | Rev: | 5'-TGTAGCTCACCTTCCACACG |
| *DES* | Fwd: | 5'-GGAAGCCGAGGAATGGTACA |
| | Rev: | 5'-TCGATCTCGCAGGTGTAGGA |
| *MYOG* | Fwd: | 5'-TGGGCGTGTAAGGTGTGTAA |
| | Rev: | 5'ACCTTCTTGAGTCTGCGCTT |
| *TNNT1* | Fwd: | 5'-CCTCTGATCCCGCCAAAGAT |
| | Rev: | 5'-GGTCCTTTTCCATGCGCTTC |

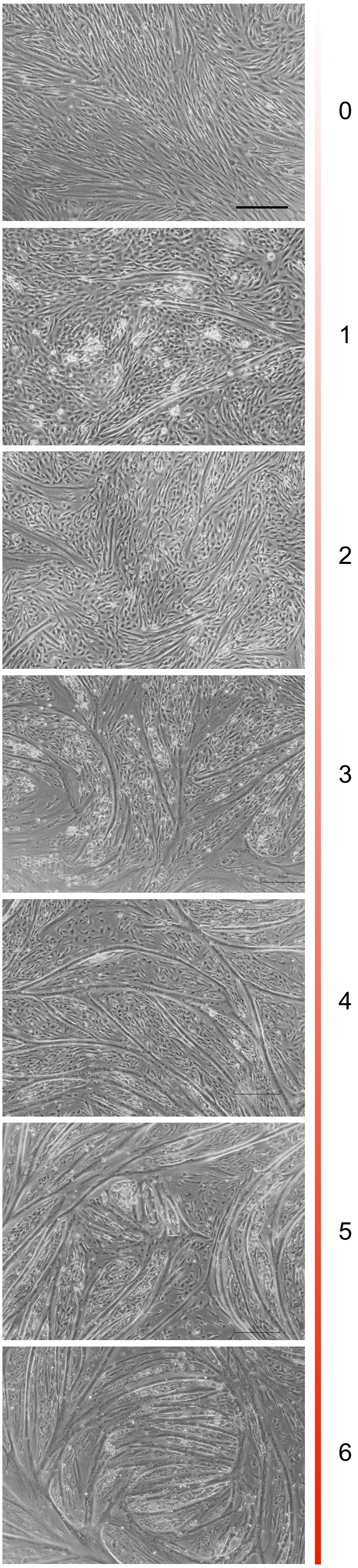


**Figure S3: Representative brightfield microscopy images corresponding to myogenic differentiation scale from 0 (no differentiation) to 6 (large myotubes, maximum extent of differentiation) used for visual assessment.**
